# Causal evidence for the involvement of Broca’s area in second language acquisition: A longitudinal HD-tDCS study

**DOI:** 10.1101/2022.12.19.520902

**Authors:** Daniel Gallagher, Kyosuke Matsumoto, Shinri Ohta

**Affiliations:** Department of Linguistics, Kyushu University, Fukuoka, Japan

**Keywords:** longitudinal HD-tDCS, inferior frontal gyrus, second language acquisition

## Abstract

A wealth of correlational evidence suggests that Broca’s area (the left inferior frontal gyrus) plays a role in second language acquisition. With the use of highly focal non-invasive brain stimulation, evidence for a targeted brain region’s causal role in some cognitive behavior can be assessed. Over three sessions, each one week apart, we used online anodal high-definition transcranial direct current stimulation (HD-tDCS) over Broca’s area during a novel foreign grammar training session. During training, participants who were naïve to Spanish were tasked with acquiring present tense conjugation rules for the three Spanish verb endings. In Session 1, we observed significant improvement in performance in two linguistic (reception and production) tasks but not in a non-linguistic (working memory) task. Sessions 2 and 3 were subject to a ceiling effect, which obscured any potentially long-lasting effects of the stimulation. We interpret these results as support for the idea that Broca’s area has languagelike specificity that is not limited to receptive or productive processes and as evidence that Broca’s area plays a causal role in foreign grammar acquisition.

## 1. Introduction

Functional neuroimaging methods, which measure neural activity via hemodynamic response, electrical signals on the scalp, magnetic fields around the scalp, etc. inherently suffer from an inability to separate correlation from causation. Counteracting the resulting dearth of causal evidence attainable by these traditional methods, transcranial electrical stimulation (tES) offers a clearer window into the workings of the mind. In this study, we employ anodal high-definition transcranial direct current stimulation (HD-tDCS) over Broca’s area during three foreign grammar training sessions to show for the first time that Broca’s area is causally linked to explicit foreign grammar acquisition.

Broca’s area (left inferior frontal gyrus (lIFG)/BA 44/45) has been implicated in various language-related tasks via a variety of neuroimaging techniques. To name a few, fMRI has indicated that the lIFG plays a role in phonetic encoding (Papoutsi et al., 2009); MEG/fMRI has shown stronger activation of Broca’s area during verb generation (Pang, Wang, Malone, Kadis, & Donner, 2011); electrocorticography has revealed a cortical network for speech planning that includes Broca’s area (Castellucci, Kovach, Howard, Greenlee, & Long, 2022), and fNIRS has shown that bilingual interpretation relies on functional connectivity between Broca’s area and Wernicke’s area (He & Hu, 2022). While these studies offer strong correlational evidence, they are, by nature, correlation studies. Thus, they fail to distinguish whether Broca’s area activation, its connecting white matter density, its blood flow, etc. cause the observed cognitive linguistic behavior or simply arise as a byproduct of or in congruence with other cognitive processes.

By providing a safe, non-invasive method for directly modulating neuronal activation, tES is invaluable for establishing causal relationships in cognitive behavioral research. In anodal tDCS experiments, electrodes placed on a participant’s scalp flow a direct current through a targeted region of the brain to increase cortical excitability and improve cognitive function (see Thair, Holloway, Newport, & Smith, 2017 for general review and Jacobson, Koslowsky, & Lavidor, 2012 for a review on polarity effects).

Generally, in HD-tDCS, fewer and smaller electrodes are used than in conventional tDCS. This setup serves the same purpose as conventional tDCS but with a more precise focality (Villamar et al., 2013), leading to a more robust localization of brain function. HD-tDCS has also been shown to have a more delayed peak and longer lasting effects compared to conventional tDCS (Kuo et al., 2013).

(HD-)tDCS has been used to explore both clinical and experimental avenues. Clinically, for example, tDCS has consistently shown promising results in the treatment of post-stroke aphasia (de Aguiar, Paolazzi, & Miceli, 2015; Elsner, Kugler, Pohl, & Mehrholz, 2013; Holland, Crinion, Holland, & Crinion, 2012). Experimentally, tDCS has robustly elicited a variety of behavioral responses. Some of the most commonly observed phenomena include the improvement of novel vocabulary acquisition (Fiori, Kunz, Kuhnke, Marangolo, & Hartwigsen, 2018; Kurmakaeva et al., 2021; Owusu & Burianová, 2020; Perikova et al., 2022), improvement in phonemic or semantic fluency (Cattaneo, Pisoni, & Papagno, 2011; Holland et al., 2011), and improvement of sentence comprehension (Giustolisi, Vergallito, Cecchetto, Varoli, & Romero Lauro, 2018; Johari, Riccardi, Malyutina, Modi, & Desai, 2021; Vergallito et al., 2020).

Most pertinent to the present study, experiments have shown an improvement of artificial grammar acquisition by using anodal tDCS over Broca’s area/lIFG (De Vries et al., 2010; Riley & Bonilha, 2021), but to our knowledge, tDCS has not yet been applied to the acquisition of natural foreign language grammar acquisition. We therefore use anodal HD-tDCS to test the hypothesis that Broca’s area is causally linked to foreign language grammar acquisition.

It is self-evident that multi-session studies provide insight into the long-lasting effects of methods like tDCS. Indeed, there has been some pushback that not enough longitudinal tDCS studies have been conducted (Arciuli & von Koss Torkildsen, 2012). On the other hand, some recent studies on working memory and aphasia shows that tDCS yields significant improvement over sham stimulation even when analyzed longitudinally (Berryhill, 2017; Fridriksson et al., 2019). Still, the shortage of non-clinical language-related longitudinal tDCS studies leaves a gap in the literature. Thus, in our study we repeat our experimental procedure over three sessions, each a week apart.

We consider four possible outcomes in this study. If none of the tasks improve with stimulation, we could deduce that Broca’s area plays little to no role in foreign grammar acquisition. In the case where both linguistic tasks improve with stimulation while the non-linguistic task does not improve, we could interpret those results as evidence that Broca’s area plays a broad languagelike-specific role in foreign grammar acquisition. If only one linguistic task improves, we could infer that Broca’s area plays a role in foreign grammar acquisition that is specific to either receptive or productive processes. Finally, in the case where all tasks improve with stimulation, it could be seen as evidence that Broca’s area plays a more domain-general role in learning.

## 2. Materials and methods

### 2.1 Participants

28 native Japanese speakers (14 male, 14 female) participated. The average age of participants was 22.06, SD = 1.57 years. All participants were naïve to Spanish. Participants were split into two groups: 14 in the active stimulation group (7 male) and 14 in the sham stimulation group (7 male). All participants had normal eyesight (including with correction) and were right-handed (M = 99.62, SD = 1.99) as measured by the Edinburgh Handedness Inventory (Oldfield, 1971). Participants were paid 1,000JPY per hour of participation. Since each session generally lasted two hours and each participant came for three sessions, the total remuneration was generally 6,000JPY.

### 2.2 Stimuli and tasks

The experiment was divided into two phases: a training phase and a mixed-task phase. Since the training phase typically finished about two minutes before stimulation ramp down, participants had a short break from training before the mixed-task phase. Task stimuli were presented visually. Excluding electrode placement, total experiment time lasted about one hour.

#### 2.2.1 Training phase

During the training phase (which was completed during online HD-tDCS), participants were introduced to Spanish subject pronouns and verb conjugation rules for the three Spanish verb types (-ar, -er, ir). Participants first watched a video of a Spanish subject pronouns chart (with corresponding Japanese translations) filling in one-by-one, with a native Spanish speaker’s voice pronouncing each word as it appeared (Figure 1A). 6 seconds elapsed between each stimulus onset, during which time participants were instructed to verbally repeat the stimulus. Thereafter and in the same manner, verb conjugation charts for each verb type were presented to the participant (Figure 1B). For the final 5 minutes of stimulation, participants were able to refer to any of the three verb conjugation charts for free review (Figure 1C). In total, the training phase lasted approximately 20 minutes.

**Figure 1.**
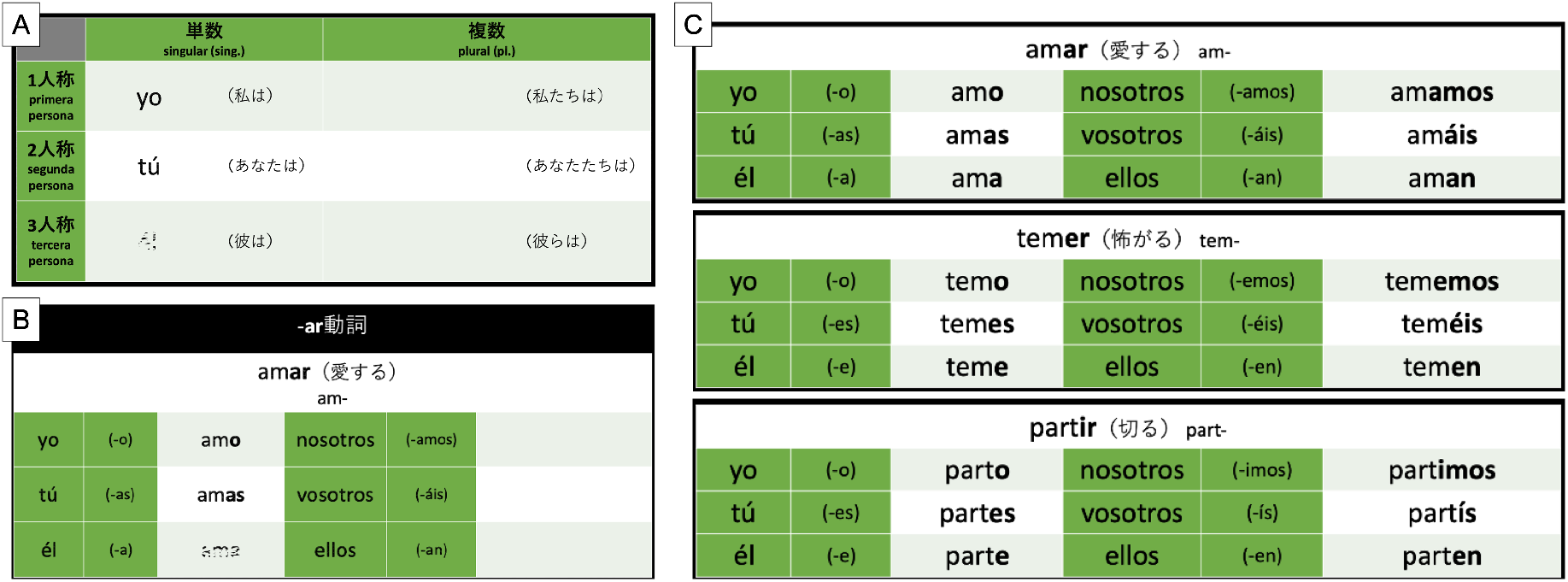
Sample screenshots of training materials. (A) A subject pronoun chart filled in one-by-one as a recording of native Spanish speaker pronouncing each word played.(B) A verb conjugation chart for Spanish -ar verbs filled in similarly to the pronoun chart in (A). The same procedure was also used to introduce -er and -ir verb conjugations. (C) For the final five minutes of stimulation, a review chart displaying conjugations of all verb types was displayed allowing participants to freely review.

#### 2.2.2 Mixed task phase

After completing the training phase, participants moved on to a mixed-task phase to assess their acquisition of the novel grammar (Figure 2). Both linguistic and non-linguistic tasks were given. Since language reception and production consist of distinctive neural subprocesses (Moreno, Rodríguez-Fornells, & Laine, 2008), for our linguistic tasks, we deemed it prudent to include both a reception and production task, in case stimulation differentially affected one or the other. The reception task was a typical grammaticality judgement task, where participants were visually presented a pronoun-verb pairing and judged whether the verb was correctly conjugated according to the given subject pronoun. The production task consisted of an unconjugated verb and a subject pronoun, and the participant was required to type the correct conjugation of the verb on a standard QWERTY keyboard. The non-linguistic task was a working memory word-recall task, where six unconjugated Spanish verbs were presented serially in quick succession for 200 ms durations. The final screen gave four options, and participants were required to select the word that was in the preceding sequence. The full mixed-task procedure took approximately 40 minutes.

**Figure 2.**
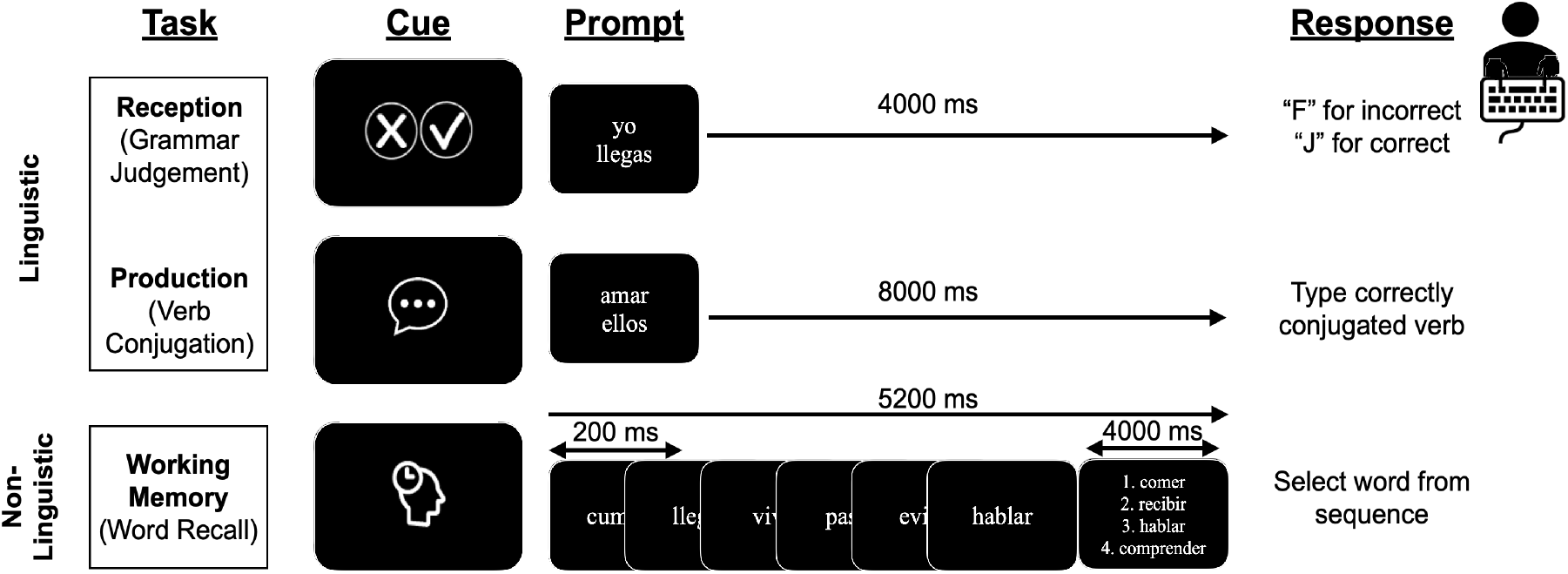
Schematic of the three stimuli and task types. Two linguistic tasks (reception and production) were compared with a control, non-linguistic (working memory) task.

This procedure of the training phase with online stimulation followed by the mixed-task phase was followed exactly across three sessions, each separated by a week.

### 2.3 Stimulation

Participants were equally divided among an active stimulation group and a sham stimulation group and underwent HD-tDCS while acquiring a novel foreign grammar in the training phase. In both groups, participants wore an electrode cap that was calibrated to their scalp by measuring the location of the Cz electrode, defined as the intersection between the midline between the inion and nasion, and the line measured from the left to right tragus. With the cap properly oriented, 5 Ag/AgCl sintered ring electrodes with an inner diameter of 5.5 mm, outer diameter of 12 mm, and thickness of 1 mm were used for stimulation and placed in a 4×1 ring geometry centered around FC5, according to the international 10-20 system for EEG recording (Figure 3A). To give anodal stimulation, the center electrode (FC5) was oriented with an anodal polarity and the surrounding electrodes (C5, FT7, F5, and FC3) with a cathodal polarity. Each cathodal electrode was set to a current of −0.25 mA, thereby giving the anodal central electrode a current of 1.0 mA. The resulting stimulation to Broca’s area is simulated in Figure 3B.

**Figure 3.**
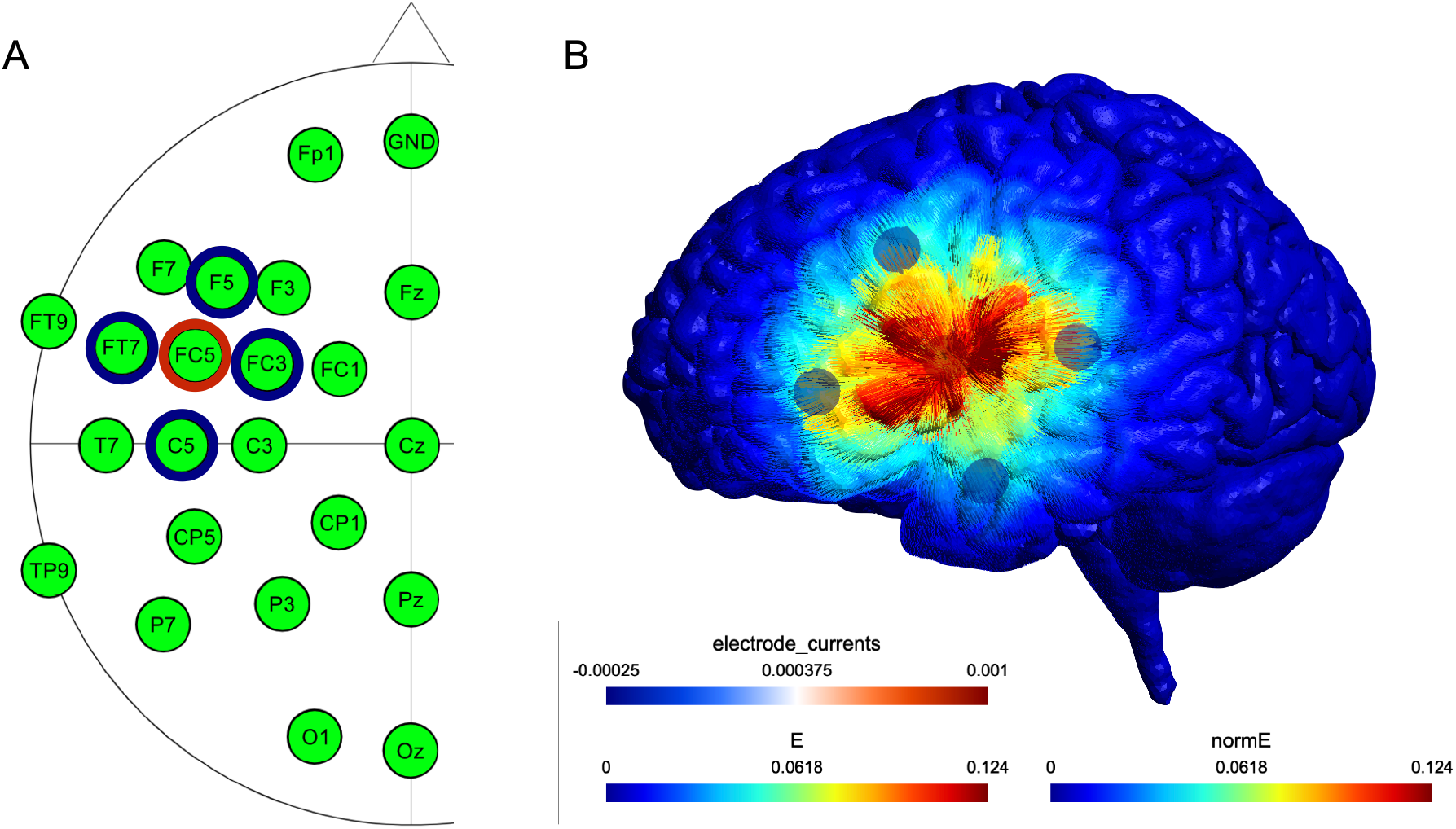
*A) Stimulation electrode configuration. Red: anodal, blue: cathodal. B) Current source simulation projected onto a canonical brain, calculated by SimNIBS (ver. 3.2.6)* (Thielscher, Antunes, & Saturnino, 2015)

### 2.4 Data Analysis

Data was analyzed using RStudio 2021.09.0. Using the “anovakun” function version 4.8.5 (Iseki, 2016), two-way analysis of variance (ANOVA) was performed to test for significant effects, defined by an alpha level of 0.05. Eta squared (η^2^) and partial eta square (η_p_^2^) were calculated to estimate effect size. We additionally ran various generalize linear mixed effects (GLME) models with binomial distribution for accuracies and linear mixed effect (LME) models for reaction time, using ANOVA to find the best-fit model for the observed data.

## 3 Results

### 3.1 Accuracies

Accuracy data was first analyzed by task, as shown in the line graphs of Figure 4. For the reception task, a significant main effect of stimulation was found (*F*(1,22) = 12.8975, *p* = 0.0016, η^2^ = 0.2383), along with a significant main effect of session (*F*(2,44) = 67.3133, *p* < 0.0001, η^2^ = 0.6419), and a significant interaction between stimulation and session (*F*(2, 44) = 13.8640,*p* < 0.0001, η^2^ = 0.2698). Post-hoc analysis showed a significant effect of stimulation during Session 1 (*F*(1,22) = 18.3946, *p* = 0.0003, η_p_^2^ = 0.4556), but not during sessions 2 or 3 (*F*(1,22) = 0.0580, *p* = 0.8119 and *F*(1,22) = 3.8930, *p* = 0.0612, respectively). Similarly in the production task, significant main effects of stimulation (*F*(1,22) = 9.8235, *p* = 0.0048, η^2^ = 0.2706) and session (*F*(2,44) = 169.6899, *p* < 0.0001, η^2^ = 0.6803) were observed along with a significant interaction between them (*F*(2,44) = 20.4570, *p* < 0.0001, η^2^ = 0.2042). Again, post-hoc analysis showed a significant effect of stimulation during Session 1 (*F*(1,22) = 19.3448, *p* = 0.0002, η_p_^2^ = 0.4679), but not during sessions 2 or 3 (*F*(1,22) = 0.0580, *p* = 0.1321 and *F*(1,22) = 3.8930, p = 0.1894, respectively). In the non-linguistic working memory task, no significant main effect of stimulation was observed (*F*(1,22) = 0.1065, *p* = 0.7472). Finally, in both groups, a learning effect was observed between sessions, indicated by horizontal bars in Figure 4.

**Figure 4.**
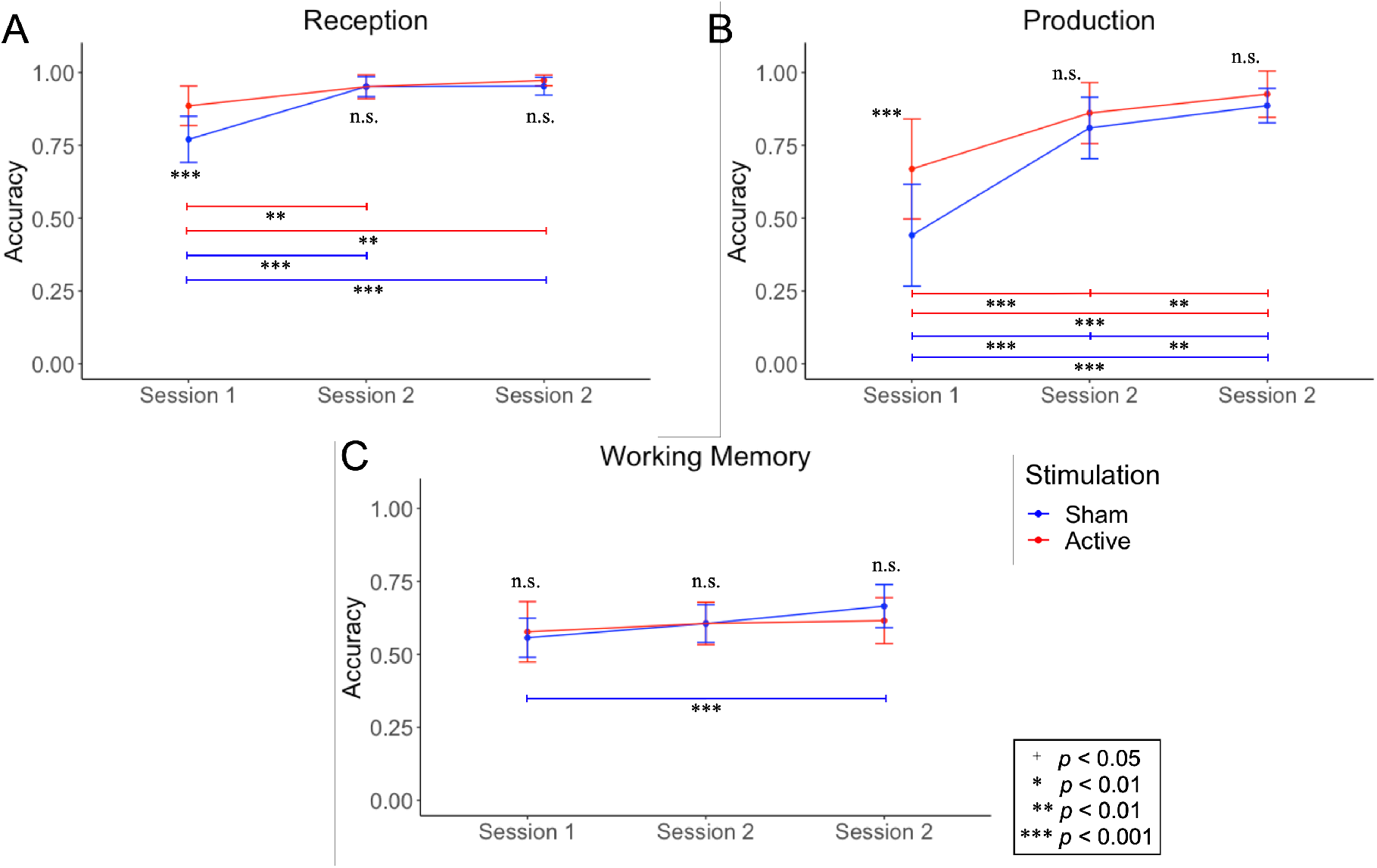
Line graphs of accuracies for each task type in each session. Active stimulation group accuracies are shown in red, sham group accuracies are shown in blue. A learning effect between sessions is depicted with color-coded horizontal bars. A) Reception task. B) Production task. C) Working memory task.

Accuracy data was subsequently analyzed by session, as shown in the bar plots of Figure 5. For Session 1 (Figure 5A), significant main effects of stimulation (*F*(1,24) = 14.2149, *p* = 0.0009, η^2^ = 0.2147) and task were found (*F*(2,48) = 52.9827, *p* < 0.0001, η^2^ = 0.5418), along with a significant interaction between stimulation and task type (*F*(2,48) = 5.9819, *p* = 0.0048, η^2^ = 0.1178). Post-hoc analysis showed a significant effect of stimulation on the two linguistic tasks, reception (*F*(1,24) = 16.0521, *p* = 0.0005, η_p_^2^ = 0.4007) and production (*F*(1,24) = 11.1500, *p* = 0.0027, η_p_^2^ = 0.3172), but not on the non-linguistic working memory task (F(1,24) = 0.3438, *p* = 0.5631). A significant main effect of task was also found for sessions 2 (Figure 5B, *F*(2,44) = 173.4412, *p* < 0.0001, η^2^ = 0.7889) and 3 (Figure 5C, F(2,44) = 233.9980, *p* < 0.0001, η^2^ = 0.8286), but no significant main effect of stimulation was observed (F(1,24) = 0.7010, *p* = 0.4107 and F(1,22) = 0.0334, *p* = 0.8567, respectively). However, in Session 3, there was a significant interaction between stimulation and task (*F*(2,44) = 4.3277, *p* = 0.0192, η^2^ = 0.0822), with post-hoc analysis showing only a marginally significant effect of stimulation in the reception task (F(1,22) = 2.8930, *p* = 0.0612, η_p_^2^ = 0.1509).

**Figure 5.**
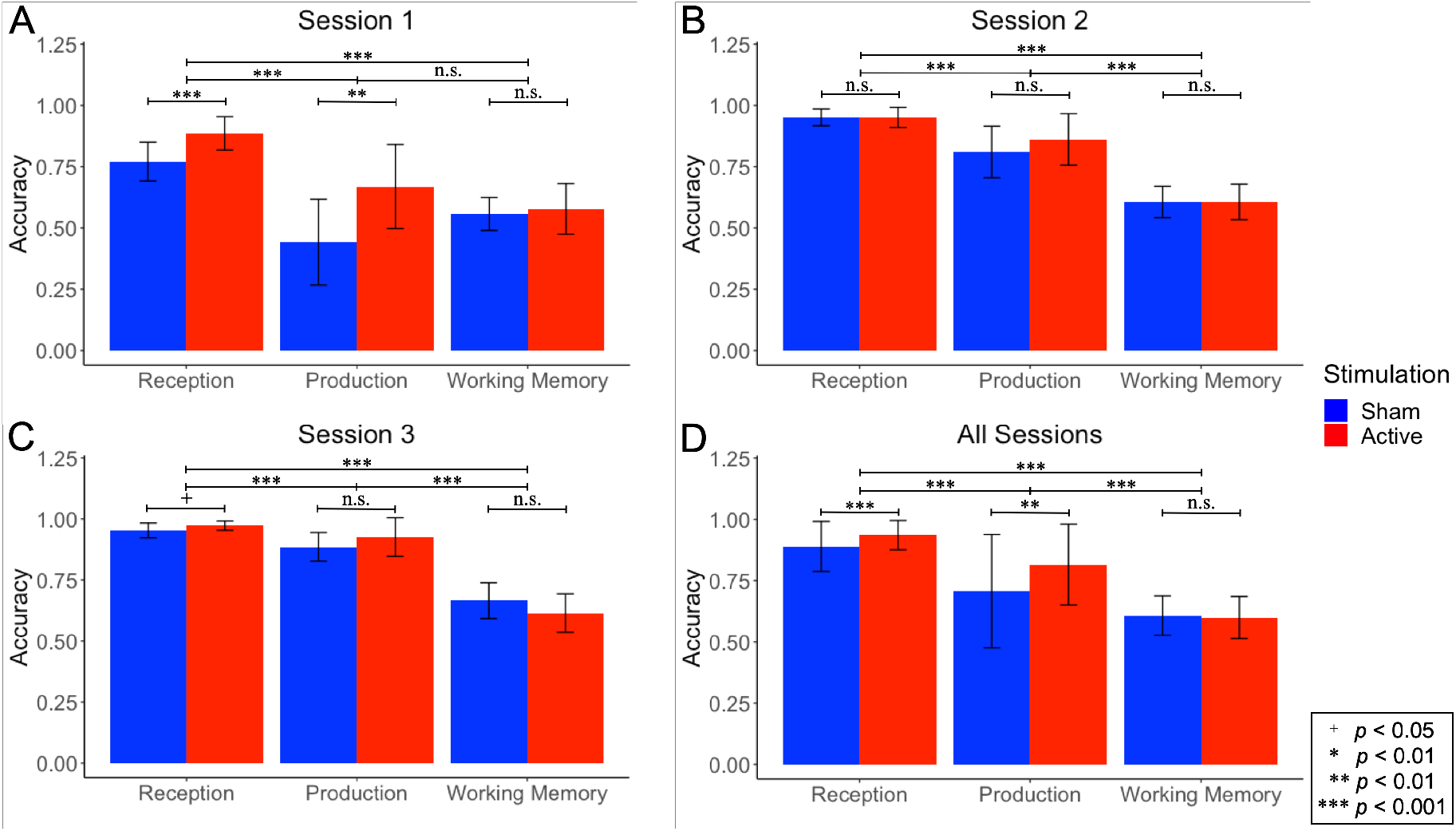
Bar plots of accuracies for all tasks across each session. Active stimulation group accuracies are shown in red, sham group accuracies are shown in blue. A) Session 1, B) Session 2, C) Session 3, D) All sessions.

For all sessions combined (Figure 5D), significant main effects of stimulation (*F*(1,24) = 14.2149, *p* = 0.0009, η^2^ = 0.1416) and task were observed (*F*(2,48) = 52.9827, *p* < 0.0001, η^2^ = 0.2629), along with a significant interaction between stimulation and task (*F*(2,48) = 5.9819, *p* = 0.0048, η^2^ = 0.0387). Post-hoc analysis of all sessions showed a significant effect of stimulation on the linguistic reception and production tasks (*F*(1,24) = 16.0521, *p* = 0.0005, η_p_^2^ = 0.4007 and *F*(1,24) = 11.1500, *p* = 0.0027, η_p_^2^ = 0.3172, respectively) but not on the non-linguistic working memory task (*F*(1,24) = 0.3438, *p* = 0.5631), which ultimately agrees with Session 1 results.

Various GLME models with binomial distribution models were also compared to find the best fit model (Table 1). We tested every combination of task (reception, production, working memory), task type (linguistic, working memory), stim (active, sham), and session (1, 2, 3) being treated as fixed effects, excluding combinations including both task and task type, since these were not independent of each other. In every model, participant was treated as a random effect. The Akaike Information Criterion (AIC), which shows the relative information value of a model, was used to calculate the best fit model, with lower AIC scores corresponding to a better fit. Using the AIC, we show that Model 10 (accuracy ~ task * stim * session + (1|participant)) is the best fit model, treating task, stimulation, and session as fixed effects and participant as a random effect.

**Table 1:** Various GLME models tested for accuracy and their corresponding Akaike Information Criterion (AIC) values.

| Model No. | GLME Model | AIC |
| --- | --- | --- |
| 1 | accuracy ~ task + (1 \| participant) | 13576 |
| 2 | accuracy ~ task type + (1 \| participant) | 13966 |
| 3 | accuracy ~ stim + (1 \| participant) | 14879 |
| 4 | accuracy ~ session + (1 \| participant) | 14434 |
| 5 | accuracy ~ task * stim + (1 \| participant) | 13517 |
| 6 | accuracy ~ task * session + (1 \| participant) | 12818 |
| 7 | accuracy ~ task type * stim + (1 \| participant) | 13908 |
| 8 | accuracy ~ task type * session + (1 \| participant) | 13245 |
| 9 | accuracy ~ stim * session + (1 \| participant) | 14398 |
| **10** | **accuracy ~ task \* stim \* session + (1 \| participant)** | **12732** |
| 11 | accuracy ~ task type * stim * session + (1 \| participant) | 13164 |

**Table 2:** Various LME models tested for RT and their corresponding AIC values.

| Model No. | LME Model | AIC |
| --- | --- | --- |
| 1 | rt ~ task + (1 \| participant) | 487.19 |
| 2 | rt ~ task type + (1 \| participant) | 883.88 |
| 3 | rt ~ stim + (1 \| participant) | 952.99 |
| 4 | rt ~ session + (1 \| participant) | 948.21 |
| 5 | rt ~ task * stim + (1 \| participant) | 487.48 |
| 6 | rt ~ task * session + (1 \| participant) | 362.69 |
| 7 | rt ~ task type * stim + (1 \| participant) | 886.19 |
| 8 | rt ~ task type * session + (1 \| participant) | 877.00 |
| 9 | rt ~ stim * session + (1 \| participant) | 952.96 |
| 10 | **rt ~ task \* stim \* session + (1 \| participant)** | **360.51** |
| 11 | rt ~ task type * stim * session + (1 \| participant) | 886.80 |

Once again, using AIC to estimate prediction error, it can be seen that the best fit LME model (Model 10: RT ∼ task * stim * session + (1|participant)) uses stimulation, task, and session as fixed effects and participant as a random effect. This is the same model as was determined to be best fit for accuracy.

### 3.2 Response times (RTs)

Response time (RT) data was analyzed in the same way as accuracy data. Figure 6 shows line graphs for each task across sessions. For the reception task, a significant main effect of stimulation was observed (*F*(1,22) = 8.0641, *p* = 0.0095, η^2^ = 0.2435) across all sessions, however post-hoc analysis revealed no significant effect of stimulation in any one of the three sessions. The production task and working memory task showed no effect of stimulation (F(1,22) = 0.7835, *p* = 0.3856 and F(1,22) = 2.2334, *p* = 0.1493, respectively). A learning effect was observed across both groups between sessions in the linguistic tasks, but not in the working memory task, as indicated by the horizontal bars in Figure 6A and 6B.

**Figure 6.**
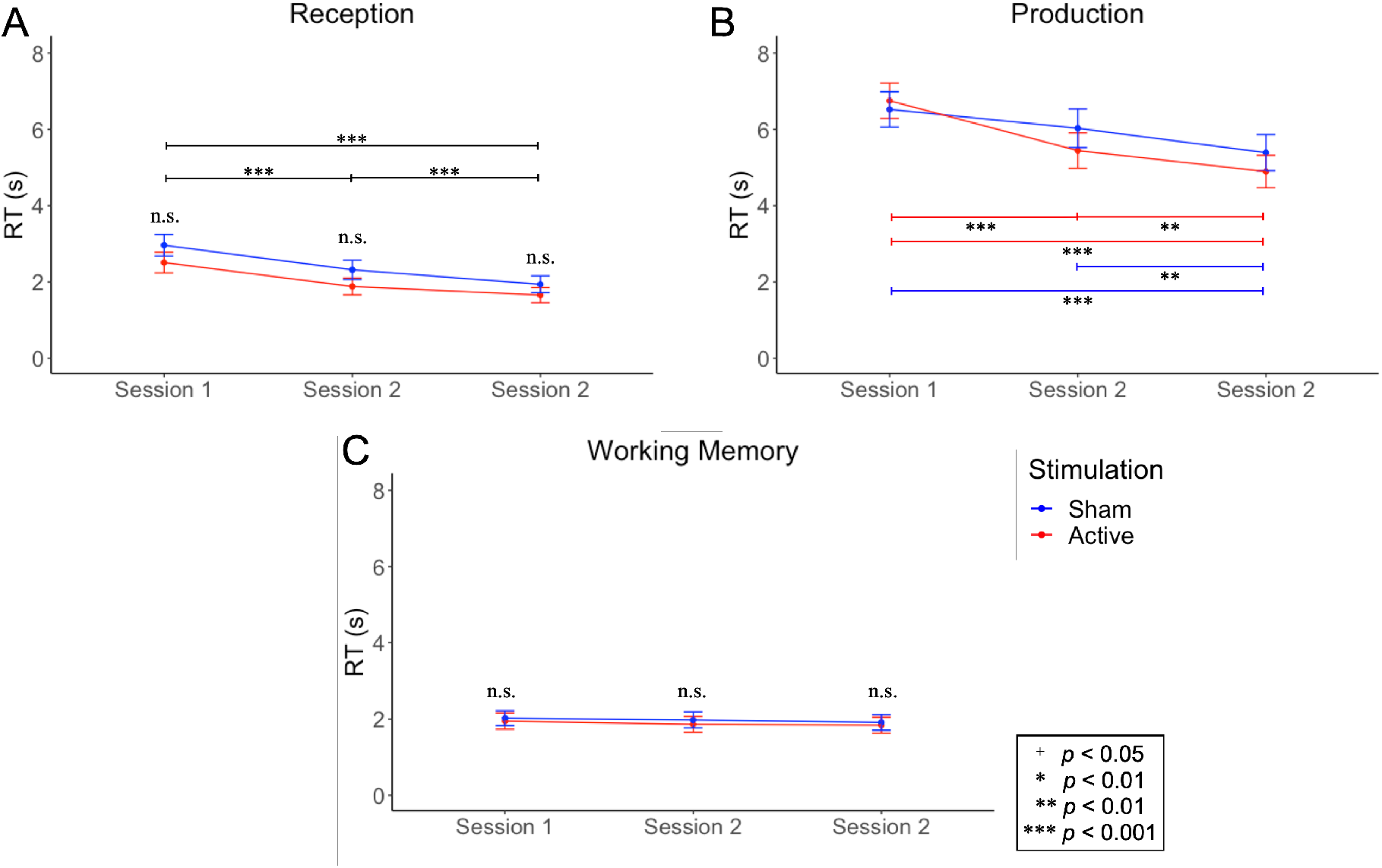
Line graphs of RT for each task type in each session. Active stimulation group RTs are shown in red, sham group RTs are shown in blue. A learning effect between sessions is depicted with color-coded horizontal bars, however since the reception task showed no significant interaction between stimulation and session, the black bar indicates a learning effect across both active and sham groups. A) Reception task. B) Production task. C) Working memory task.

A bar plot is shown in Figure 7 to compare RTs of the different tasks within each session. In Session 1, no significant main effect of stimulation was found (*F*(1,23) = 0.4612, *p* = 0.5036), but a significant main effect of task was found (*F*(2,48) = 2.1146, *p* < 0.0001, η^2^ = 0.9090). There was no significant interaction between stimulation and task (*F*(2,48) = 2.1146, *p* = 0.1318). In Session 2, there were significant main effects of stimulation and task (*F*(1,24) = 5.5136, *p* = 0.0274, η^2^ = 0.1035 and *F*(2,48) = 449.5468, *p* < 0.0001, η^2^ = 0.9049, respectively), but there was no interaction between them (*F*(2,48) = 1.5428, *p* = 0.2242). In Session 3, there was a marginally significant main effect of stimulation (*F*(1,22) = 3.3820, *p* = 0.0795, η^2^ = 0.0867), there was a significant main effect of task (*F*(2,44) = 717.0930, *p* < 0.0001, η^2^ = 0.9264), and there was no interaction between stimulation and task (*F*(2,44) = 1.9955, *p* = 0.1481). For all sessions, there was a marginally significant main effect of stimulation (*F*(1,24) = 3.1449, *p* = 0.0889, η^2^ =0.0625), there was a significant main effect of task (*F*(2,48) = 730.0933, *p* < 0.0001, η^2^ =0.9289), and there was no significant interaction between stimulation and task (*F*(2,48) = 0.8661, *p* = 0.1481).

**Figure 7.**
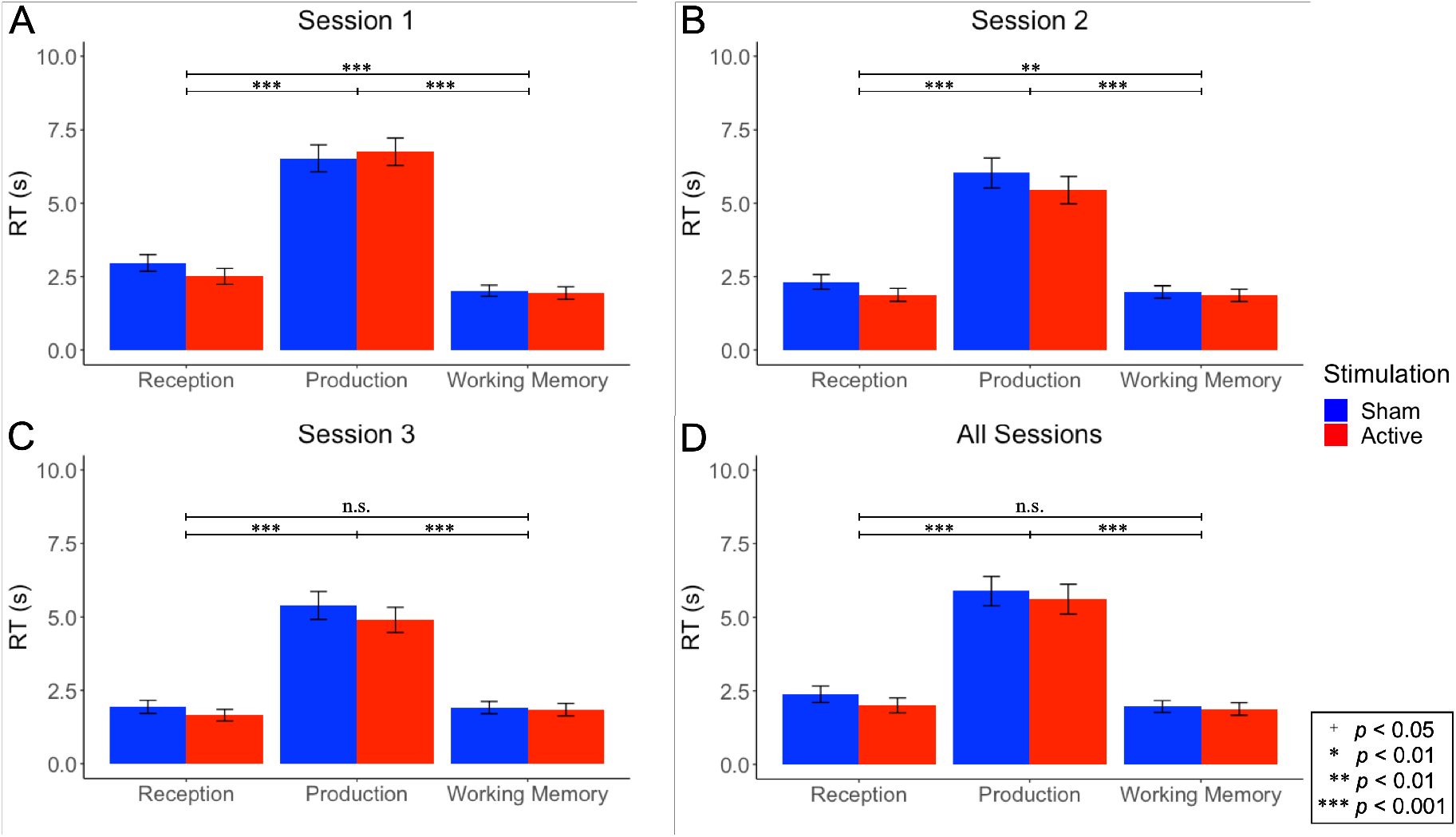
Bar plots of RTs for all tasks across each session. Active stimulation group RTs are shown in red, sham group RTs are shown in blue. A) Session 1, B) Session 2, C) Session 3, D) All sessions.

The same models tested for accuracy (Table 1) were tested for RT, using LME instead of GLME with binomial distribution, and in the same way were compared to find the best fit model.

## 4 Discussion

The first key finding of our experiment is that in Session 1, the active stimulation group performed significantly better than sham at applying their newly acquired Spanish conjugation rules. Importantly, stimulation of Broca’s area differentially affected linguistic task performance, having no significant effect on a non-linguistic working memory task. Specifically, we observed cognitive improvement most clearly in the accuracy results (Figures 4 and 5), with reception task accuracy improving by 11.53% and production task accuracy by 22.73%. Working memory task accuracy rose by only 2.05% and was nonsignificant.

When looking at all sessions combined, the active stimulation group significantly outperformed the sham stimulation group (Figure 5) in task accuracy. However, because neither Session 2 nor Session 3 showed significant differences between groups, the combined results simply showcase the strength of the effect observed in Session 1.

Since the training phase consisted of explicit grammar instruction, it was not surprising that both groups demonstrated a learning effect (Figures 4 and 6), irrespective of stimulation type. While we hoped to observe whether active stimulation had long-term effects on grammar acquisition, the magnitude of the learning effect ultimately led to a ceiling effect starting with Session 2. This meant that participants in both groups performed at high enough accuracies to minimize or altogether erase any intergroup differences in accuracy due to stimulation. Still, in Session 3, the active stimulation group performed marginally significantly better on the reception task, suggesting sustained long-term effects of stimulation. Nevertheless, in order to address the question of long-term effects of stimulation more adequately, a more difficult and/or complex task would be required.

As for RTs, the active stimulation group responded significantly faster than sham in the reception task when analyzing all sessions in the production task. However, no individual session rose to the level of significance. Since participants were not instructed to answer quickly and tasks involved applying a newly acquired grammar, it is reasonable to assume that participants in both groups took great care when answering, paying little mind to the time elapsed while responding.

Given that stimulation over Broca’s area improved both reception and production task performance but not working memory task performance, we can surmise that the underlying linguistic processing being carried out by Broca’s area has a broad languagelike specificity, not limited to either receptive or productive processes. On the other hand, since task was shown to be a better predictor than the broader category of task type, stimulation may affect some language tasks more strongly than others. Alternatively, the baseline difficulty of the task itself (irrespective of stimulation) may modulate the magnitude of the stimulation effect. Regardless, we have ultimately shown that online anodal tDCS over Broca’s area differentially improves performance on linguistic tasks related to a newly learned natural foreign language grammar, thus providing strong evidence for a causal relationship between Broca’s area and foreign language grammar acquisition.

As previously mentioned, the tasks used herein were easy enough for the participants to induce a ceiling effect by the second session, thereby limiting the usefulness of the longitudinal analysis. To that end, future studies may find more suitable behavioral tasks for longitudinal analysis.

Finally, tDCS provides no data of the neural processing of a given stimulus. Therefore, a co-registration study employing electroencephalography alongside tDCS would be fitting to clarify the neurophysiological effects induced by tDCS.

After using online anodal HD-tDCS over Broca’s area during a novel foreign grammar acquisition training session, we observed a significant and differential improvement in performance on tasks related to the novel grammar, thereby providing evidence for a causal relationship between Broca’s area and foreign language acquisition. While these results may be unsurprising in light of the wealth of correlational evidence provided by previous studies, we believe that the establishment of the causal role Broca’s area plays in natural language acquisition is a necessary and critical step in the broader goal of elucidating the neurophysiology of language processing.

## References

Arciuli, J., & von Koss Torkildsen, J. (2012). Advancing our understanding of the link between statistical learning and language acquisition: The need for longitudinal data. Frontiers in Psychology, 3(AUG), 1–9. https://doi.org/10.3389/fpsyg.2012.00324

Berryhill, M. E. (2017). Longitudinal tDCS: Consistency across working memory training studies. AIMS Neuroscience, 4(2), 71–86. https://doi.org/10.3934/Neuroscience.2017.2.71

Castellucci, G. A., Kovach, C. K., Howard, M. A., Greenlee, J. D. W., & Long, M. A. (2022). A speech planning network for interactive language use. Nature, 602(7895), 117–122. https://doi.org/10.1038/s41586-021-04270-z

Cattaneo, Z., Pisoni, A., & Papagno, C. (2011). Transcranial direct current stimulation over Broca’s region improves phonemic and semantic fluency in healthy individuals. Neuroscience, 183, 64–70. https://doi.org/10.1016/j.neuroscience.2011.03.058

de Aguiar, V., Paolazzi, C. L., & Miceli, G. (2015). tDCS in post-stroke aphasia: The role of stimulation parameters, behavioral treatment andpatient characteristics. Cortex, 63, 296–316. https://doi.org/10.1016/j.cortex.2014.08.015

De Vries, M. H., Barth, A. C. R., Maiworm, S., Knecht, S., Zwitserlood, P., & Flöel, A. (2010). Electrical stimulation of Broca’s area enhances implicit learning of an artificial grammar. Journal of Cognitive Neuroscience, 22(11), 2427–2436. https://doi.org/10.1162/jocn.2009.21385

Elsner, B., Kugler, J., Pohl, M., & Mehrholz, J. (2013). Transcranial direct current stimulation (tDCS) for improving function and activities of daily living in patients a er stroke (Review). Cochrane Database of Systematic Reviews, (11). https://doi.org/10.1002/14651858.CD009645.pub2.www.cochranelibrary.com

Fiori, V., Kunz, L., Kuhnke, P., Marangolo, P., & Hartwigsen, G. (2018). Transcranial direct current stimulation (tDCS) facilitates verb learning by altering effective connectivity in the healthy brain. NeuroImage, 181(July), 550–559. https://doi.org/10.1016/j.neuroimage.2018.07.040

Fridriksson, J., Basilakos, A., Stark, B. C., Rorden, C., Elm, J., Gottfried, M., … Bonilha, L. (2019). Transcranial direct current stimulation to treat aphasia: Longitudinal analysis of a randomized controlled trial. Brain Stimulation, 12(1), 190–191. https://doi.org/10.1016/j.brs.2018.09.016

Giustolisi, B., Vergallito, A., Cecchetto, C., Varoli, E., & Romero Lauro, L. J. (2018). Anodal transcranial direct current stimulation over left inferior frontal gyrus enhances sentence comprehension. Brain and Language, 176(October 2017), 36–41. https://doi.org/10.1016/j.bandl.2017.11.001

He, Y., & Hu, Y. (2022). Functional Connectivity Signatures Underlying Simultaneous Language Translation in Interpreters and Non-Interpreters of Mandarin and English: An fNIRS Study. Brain Sciences, 12(2). https://doi.org/10.3390/brainsci12020273

Holland, R., Crinion, J., Holland, R., & Crinion, J. (2012). Can tDCS enhance treatment of aphasia after stroke ? Can tDCS enhance treatment of aphasia after stroke ?, 7038. https://doi.org/10.1080/02687038.2011.616925

Holland, R., Leff, A. P., Josephs, O., Galea, J. M., Desikan, M., Price, C. J., … Crinion, J. (2011). Speech facilitation by left inferior frontal cortex stimulation. Current Biology, 21(16), 1403–1407. https://doi.org/10.1016/j.cub.2011.07.021

Iseki, R. (2016). Anovakun (Version 4.8.5).

Jacobson, L., Koslowsky, M., & Lavidor, M. (2012). tDCS polarity effects in motor and cognitive domains: a meta-analytical review. Experimental Brain Research, 216, 1–10. https://doi.org/10.1007/s00221-011-2891-9

Johari, K., Riccardi, N., Malyutina, S., Modi, M., & Desai, R. H. (2021). HD-tDCS over motor cortex facilitates figurative and literal action sentence processing. Neuropsychologia, 159(July), 107955. https://doi.org/10.1016/j.neuropsychologia.2021.107955

Kuo, H. I., Bikson, M., Datta, A., Minhas, P., Paulus, W., Kuo, M. F., & Nitsche, M. A. (2013). Comparing cortical plasticity induced by conventional and high-definition 4 × 1 ring tDCS: A neurophysiological study. Brain Stimulation, 6(4), 644–648. https://doi.org/10.1016/j.brs.2012.09.010

Kurmakaeva, D., Blagovechtchenski, E., Gnedykh, D., Mkrtychian, N., Kostromina, S., & Shtyrov, Y. (2021). Acquisition of concrete and abstract words is modulated by tDCS of Wernicke’s area. Scientific Reports, 11(1), 1–12. https://doi.org/10.1038/s41598-020-79967-8

Moreno, E. M., Rodríguez-Fornells, A., & Laine, M. (2008). Event-related potentials (ERPs) in the study of bilingual language processing. Journal of Neurolinguistics, 21(6), 477–508. https://doi.org/10.1016/j.jneuroling.2008.01.003

Oldfield, R. C. (1971). The Assessment and Analysis of Handedness: The Edinburgh Inventory. Neuropsychologia, 9, 97–113.

Owusu, B. K., & Burianová, H. (2020). Transcranial direct current stimulation improves novel word recall in healthy adults. Journal of Neurolinguistics, 53(March 2019), 100862. https://doi.org/10.1016/j.jneuroling.2019.100862

Pang, E. W., Wang, F., Malone, M., Kadis, D. S., & Donner, E. J. (2011). Localization of Broca’s area using verb generation tasks in the MEG: Validation against fMRI. Neuroscience Letters, 490(3), 215–219. https://doi.org/10.1016/j.neulet.2010.12.055

Papoutsi, M., De Zwart, J. A., Jansma, J. M., Pickering, M. J., Bednar, J. A., & Horwitz, B. (2009). From phonemes to articulatory codes: An fMRI study of the role of broca’s area in speech production. Cerebral Cortex, 19(9), 2156–2165. https://doi.org/10.1093/cercor/bhn239

Perikova, E., Blagovechtchenski, E., Filippova, M., Shcherbakova, O., Kirsanov, A., & Shtyrov, Y. (2022). Anodal tDCS over Broca’s area improves fast mapping and explicit encoding of novel vocabulary. Neuropsychologia, 108156. https://doi.org/10.1016/j.neuropsychologia.2022.108156

Riley, E., & Bonilha, L. (2021). TDCS-mediated artificial grammar training shows potential for improving attention and facilitating language learning in persons with aphasia. Brain Stimulation, 14(6), 1675. https://doi.org/10.1016/j.brs.2021.10.277

Thair, H., Holloway, A. L., Newport, R., & Smith, A. D. (2017). Transcranial direct current stimulation (tDCS): A Beginner’s guide for design and implementation. Frontiers in Neuroscience, 11(NOV). https://doi.org/10.3389/fnins.2017.00641

Thielscher, A., Antunes, A., & Saturnino, G. B. (2015). Field modeling for transcranial magnetic stimulation: A useful tool to understand the physiological effects of TMS? Proceedings of the Annual International Conference of the IEEE Engineering in Medicine and Biology Society, EMBS, 2015-Novem, 222–225. https://doi.org/10.1109/EMBC.2015.7318340

Vergallito, A., Varoli, E., Giustolisi, B., Cecchetto, C., Del Mauro, L., & Romero Lauro, L. J. (2020). Mind the stimulation site: Enhancing and diminishing sentence comprehension with anodal tDCS. Brain and Language, 204(November 2019), 104757. https://doi.org/10.1016/j.bandl.2020.104757

Villamar, M. F., Volz, M. S., Bikson, M., Datta, A., Dasilva, A. F., & Fregni, F. (2013). Technique and Considerations in the Use of 4×1 Ring High-definition Transcranial Direct Current Stimulation (HD-tDCS). Journal of Visualized Experiments, 77(July), 1–15. https://doi.org/10.3791/50309

